# The temporal evolution of conceptual object representations revealed through models of behavior, semantics and deep neural networks

**DOI:** 10.1101/223990

**Authors:** B. B. Bankson, M.N. Hebart, I.I.A. Groen, C.I. Baker

**Affiliations:** Section on Learning and Plasticity, Laboratory of Brain and Cognition, National Institute of Mental Health, National Institutes of Health, Bethesda, MD 20892, USA

## Abstract

Visual object representations are commonly thought to emerge rapidly, yet it has remained unclear to what extent early brain responses reflect purely low-level visual features of these objects and how strongly those features contribute to later categorical or conceptual representations. Here, we aimed to estimate a lower temporal bound for the emergence of conceptual representations by defining two criteria that characterize such representations: 1) conceptual object representations should generalize across different exemplars of the same object, and 2) these representations should reflect high-level behavioral judgments. To test these criteria, we compared magnetoencephalography (MEG) recordings between two groups of participants (*n* = 16 per group) exposed to different exemplar images of the same object concepts. Further, we disentangled low-level from high-level MEG responses by estimating the unique and shared contribution of models of behavioral judgments, semantics, and different layers of deep neural networks of visual object processing. We find that 1) both generalization across exemplars as well as generalization of object-related signals across time increase after 150 ms, peaking around 230 ms; 2) behavioral judgments explain the most unique variance in the response after 150 ms. Collectively, these results suggest a lower bound for the emergence of conceptual object representations around 150 ms following stimulus onset.

## Introduction

There is enormous variability in the visual appearance of objects, yet we can rapidly recognize them without effort, even under difficult viewing conditions (DiCarlo & Cox, 2007; Potter et al., 2013). Evidence from neurophysiological studies in human suggests the emergence of visual object representations within the first 150 ms of visual processing (Thorpe et al., 1996; Carlson et al., 2013, Cichy et al., 2014). For example, the specific identity of objects can be decoded from the magnetoencephalography (MEG) signal with high accuracy around 100 ms (Cichy et al., 2014). However, knowing when discriminative information about visual objects is available does not inform us about the nature of those representations, in particular whether they primarily reflect (low-level) visual features or (high-level) conceptual aspects of the objects (Clarke et al., 2015). To address this issue, in this study we employed multivariate MEG decoding and model-based representational similarity analysis (RSA) to elucidate the nature of object representations over time.

Previous studies have demonstrated increasing category specificity (van de Nieuwenhuijzen et al., 2013; Cichy et al., 2014), tolerance for position and size (Isik et al., 2014) and semantic information (Clarke et al., 2013) over the first 200ms following stimulus onset, suggesting some degree of abstraction from low-level visual features. However, identifying the nature of object representations is an inherently difficult problem: low-level features may be predictive of object identity, making it hard to disentangle the relative contribution of low and high-level properties to measured brain signals (Groen et al., 2017). In this study, we addressed this problem by combining tests for the generalization of object representations with methods to separate the independent contributions of low- and high-level properties. We focused on two specific criteria that would need to be fulfilled for a representation to be considered conceptual. First, a conceptual representation should generalize beyond the specific exemplar presented, not just variations of the same exemplar. Second, a conceptual representation should also reflect high-level behavioral judgments about objects (Clarke & Tyler, 2015; Wardle et al., 2016). We consider fulfillment of these two properties to provide a lower bound at which a representation could be considered conceptual.

We collected MEG and behavioral data from 32 participants allowing us to probe the temporal dynamics of conceptual object representations according to the two criteria above. To test for generalization across specific exemplars, we assessed the reliability of object representations across two independent sets of objects. Further, we assessed the relation of those object representations to behavior by comparing participants’ behavioral judgments with the MEG response patterns using RSA. Importantly, to isolate the relative contributions of low-level and conceptual properties to those MEG responses, we identified the variance uniquely explained by behavioral judgments, isolating low-level representations using early layers of a deep neural network, which have been shown to capture low-to mid-level responses in fMRI and monkey ventral visual cortex (Cadieu et al., 2014; Cichy et al., 2016a; Eickenberg & Thirion, 2017; Güçlü & van Gerven, 2015; Khaligh-Razavi & Kriegeskorte, 2014; Yamins et al., 2014; Wen et al., 2017). Finally, to achieve a more interpretable understanding of the contribution of behavior to MEG responses, we identified the unique and shared variance explained in the MEG response by behavior and two high-level conceptual models, one perceptual (upper layers in a deep neural network) and one semantic (based on word co-occurrence statistics).

## Methods

### Participants

32 healthy participants (18 female, mean 25.8, range 19-47) with normal or corrected-to-normal vision took part in this study. As a part of a pilot experiment used for purely illustrative purposes (see Figure 4a), 8 participants (5 overlap) completed the same behavioral task with a different set of stimuli. All participants gave written informed consent prior to participation in the study as a part of the study protocol (93-M-0170, NCT00001360). The study was approved by the Institutional Review Board of the National Institutes of Health and was conducted according to the Declaration of Helsinki.

### Stimuli

We created two independent sets of 84 object images each that were cropped and placed on a grey background. Each stimulus set contained a unique exemplar for each of the 84 object concepts, as shown in Figure 1a. We selected object concepts by using a combination of two word databases, one of word frequency (Corpus of Contemporary American English, Davies, 2008) and the other of word concreteness (Brysbaert et al., 2014). First, based on our corpus we selected the 5000 most frequent nouns in American English. From this set of words, we then selected nouns with concreteness ratings > 4/5. Finally, for words that would be difficult or impossible to distinguish when presented as an image (e.g. ‘woman’, ‘mother’, ‘wife’), we used only the most frequent entry. This selection left us with a set of 112 objects.

**Figure 1.**
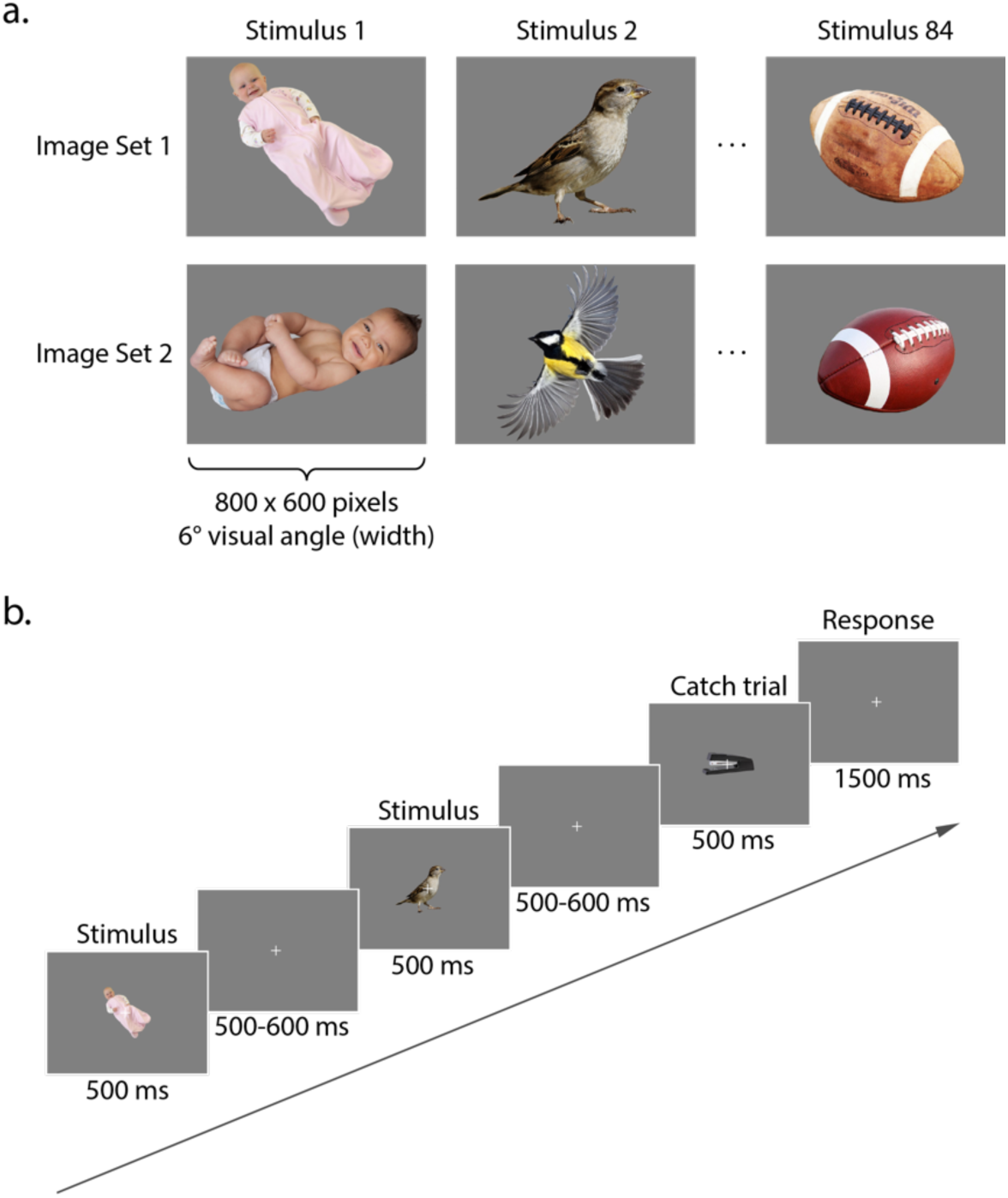
Stimulus format and trial progression. **a.** Two unique object exemplars were selected for each of the 84 object concepts used in the study. **b.** Stimuli were presented on a grey background for 500 ms, followed by fixation for 500-600 ms (catch trials: 1500 ms). All 84 stimuli from both image sets are shown in Supplemental Figure S1.

To evaluate whether those categories would be labeled consistently, we generated three distinct images of each object concept and asked three individuals who were not involved in the study to provide a verbal label for each of the three versions of the 112 objects. Images that were not labeled correctly by all raters were discarded, leaving us with 84 object concepts. From the three sets of object images, we then randomly sampled two per object concept. This generated two sets of unique object exemplars for 84 object concepts, divided into Image Set 1 and Image Set 2. The two sets of object stimuli are shown in Supplemental Figure S1.

### Procedure

#### MEG

During MEG recordings, participants were seated upright in an electromagnetically shielded MEG chamber. Stimuli were presented using the Psychophysics Toolbox (Brainard, 1997) in MATLAB (version 2016a, Mathworks, Natick, MA). Visual stimulation was controlled by a Panasonic PT-D3500U DLP projector with an ET-DLE400 lens, located outside of the chamber and projected through a waveguide and series of mirrors onto a back-projection screen in front of the participant. Participants were assigned to one of two groups and completed the experiment with either Image Set 1 or Image Set 2. All stimuli were presented on a grey background with a white fixation cross in the center (viewing distance: 70 cm, stimulus width: 6° of visual angle). Participants completed an oddball detection task, pressing a button in response to catch trials containing the oddball stimulus (desk stapler) that appeared pseudorandomly every 2-6 trials (average 4, flat distribution). On each trial (Figure 1b), an object stimulus was presented at fixation for 500 ms, followed by a variable fixation period (regular trials: pseudorandomly 500-600 ms, catch trials: 1500 ms). In addition, participants were instructed to blink their eyes only as they pressed the button of the MEG-compatible button box during catch trials, in order to avoid any eye blink artifacts at other points of the experiment. Participants completed 18 runs that were divided into 6 blocks of 3 runs each, with self-paced breaks between each block. Each run lasted 240 s, resulting in a total experimental time of 72 min. In total, participants viewed each of the 84 images 36 times over the course of the experiment.

#### Behavior: Object arrangement task

Within two days of completing the MEG session, participants took part in a follow-up behavioral experiment to provide us with behavioral estimates of the representational similarity between all possible object pairs. This was done using the object arrangement method (Goldstone 1994; Kriegeskorte & Mur, 2012). In this method, participants arrange objects in a 2D “arena” based on their subjective similarity, and the distance between the items is used to generate (n × n-1)/2 pairwise distance estimates between object pairs. Participants were seated at a distance of approximately 57 cm in front of a 30” monitor (resolution: 1440 × 900 pixels) and completed the object arrangement task on the same 84 object images used in the MEG experiment. All items were presented simultaneously but in random order and with equal distance around the circular arena (image width: 1.5° of visual angle). Participants were instructed to use the computer mouse and arrange the items according to their similarity at their own pace, taking ~20 minutes on average to complete the task. In contrast to the original implementation of this method that used additional trials with selective subsets of objects (Kriegeskorte et al., 2012), we only chose a single arrangement, based on our experience with the multi-arrangement task exhibiting very high correlations between results of the first and the last trial (unpublished data). We deliberately did not provide participants with an explicit strategy or instructions on what object features to focus, so as to not bias them to focus on any specific aspect of the stimuli. To facilitate the task, when a participant clicked on a certain image around the arena, an enlarged version spanning 150 × 200 pixels (6.75 × 9° of visual angle) was displayed in the top right of the computer screen. After completion of the experiment, we extracted the pixel-wise distance between each pair of items, yielding an 84 × 84 distance matrix for each participant. Note that the distance matrix discards the absolute position of objects and only retains their relative location, which should minimize bias related to the initial placement of objects.

#### MEG acquisition and preprocessing

MEG data were recorded continuously at a sampling rate of 1200 Hz with a 275-channel CTF whole-head MEG system (MEG International Services, Ltd., Coquitlam, BC, Canada). All analyses were conducted in MATLAB (version 2016a, The Mathworks, Natick, MA). Preprocessing was carried out using Brainstorm 3.4 (version 02/2016, Tadel et al., 2011) and custom-written code, using similar preprocessing steps as previously published MEG decoding work (Cichy et al., 2014; Grootswagers et al., 2016, Hebart et al., 2018). Recordings were available from 272 channels (dead channels: MLF25, MRF43, MRO13). The whole-head array consists of radial first-order gradiometer channels equipped with synthetic third-gradient balancing to remove background noise online. At the beginning of the experiment and after every third experimental run, participants’ head position was localized based on fiducial coil placement at the nasion, left and right preauricular points. Head position was recorded to provide the experimenter with feedback about the head position to reposition the participant’s head in the dewar if necessary. Data were bandpass filtered between 0.1 and 300 Hz, and bandstop filtered at 60 Hz and harmonics. We segmented the data into single trial bins, with each trial consisting of 100 ms baseline for normalization purposes and 1000 ms post-stimulus activity, yielding a total of 1321 time samples for each trial. Catch trials were discarded.

Three pre-analysis steps allowed us to increase SNR and reduce computational demand: PCA dimensionality reduction, temporal smoothing on PCA components, and data downsampling. Principal components analysis (PCA) was run to reduce the number of channels into the set of most descriptive components. All data for an MEG channel across trials were concatenated for PCA, and the components explaining the least variance were removed to speed-up further processing, with a maximum removal of 50 % of the components (i.e. 136 components) or 1 % of the variance, whichever was reached first (Hebart et al., 2018). Since for all participants the smallest 136 components explained less than 1 % of the variance, the data for further analyses contained 136 components. Data across all time points were normalized according to the baseline period of −100 to 0 ms relative to stimulus presentation. To do so, the mean and standard deviation of the baseline period for each component were computed, and the mean was subtracted from the data before dividing by the standard deviation. We then used a Gaussian kernel of ± 15 ms half duration at half maximum (HDHM) to temporally smooth the remaining components, and downsampled the components to 120 Hz (132 samples / trial).

### Multivariate decoding and temporal generalization analysis

#### Multivariate MEG decoding

Our goal was to study the representational dynamics during visual object recognition and the emergence of generalizable, conceptual object representations over time. To determine the amount of object information contained in the MEG signal over time, we ran time-resolved multivariate decoding of MEG data using a linear support vector machine classifier (SVM; Chang & Lin, 2011). The analysis steps were chosen according to general recommendations (Grootswagers et al., 2016) and a recent study from our lab (Hebart et al., 2018). Multivariate analyses were conducted using functions from The Decoding Toolbox (Hebart et al., 2015) and custom-written code. The following analysis steps were applied to all participants, regardless of experimental group.

First, we created supertrials by averaging 6 trials of the same object concept drawn randomly without replacement (Isik et al., 2014). For each time point, preprocessed MEG data within each supertrial were arranged as *P* dimensional measurement vectors (corresponding to the number of components from PCA preprocessing), yielding *K* pattern vectors for each time point and object concept. For each pair of object concepts and each time point, we then trained the classifier on *K-1* pattern vectors and tested it on the pair of left-out pattern vectors, yielding a decoding accuracy for each pair of object categories at each time point. Note that while leave-one-out cross-validation can lead to some overfitting to the data at hand, when the purpose is to demonstrate a statistical dependence in combination with classical statistics this is a valid approach (Hebart & Baker, 2017). The assignment to training and testing sets and resulting classification procedure was repeated 100 times for each pair of object concepts and each time point, with a new random generation of supertrials in each iteration. The resulting decoding accuracies were averaged across the 100 iterations and presented as an 84 × 84 matrix at every time point, with rows and columns indexed according to object conditions, and with the diagonal undefined. We used these matrices to evaluate average decoding accuracy at each time point by computing the average of the lower triangular matrix.

Significance for the decoding analysis was assessed using a sign permutation test. A null distribution of group means was generated by running the decoding procedure 1,000 times, randomly generating a sign-permuted accuracy per participant and averaging those values. *P*-values were determined as one minus the percentile of the original group mean in this null distribution. Those *p*-values were corrected according to the false-discovery rate (FDR) and were deemed significant if the corrected *p*-value did not exceed 0.05 (i.e. the test was one-sided).

#### Temporal generalization of object representation

While time-resolved multivariate decoding can reveal when specific mental representations are present in patterns of neural activity, it cannot identify how said patterns at one time point relate to other time points. We were interested in investigating the extent to which object-related information is static or dynamic over time, which can give us an index of how rapidly neural signals evolve. To investigate this, we conducted a cross-classification analysis over time, also known as the temporal generalization method (King & Dehaene, 2014; Meyers et al., 2008). If a classifier can successfully generalize from one time point to another, this shows that representational content is highly similar between these two time points. Conversely, if the classifier does not generalize, this shows that patterns of neural activity have evolved to an extent that representational content is no longer similar.

To carry out this temporal generalization analysis, we used the same classification approach described above; however, instead of only testing the classifier at the same time point we also tested its performance at all other time points. We repeated the analysis with all time points each serving as training data once for the classifier, and generated a 132 x 132 time-time decoding matrix that shows the extent to which our classifier generalizes across time.

#### Representational similarity analysis (RSA)

RSA is a method to analyze and compare data patterns, for example brain activity patterns with behavioral judgments or computational models (Kriegeskorte et al., 2008). Instead of comparing these patterns directly, in RSA patterns are converted to representational similarity matrices (RSMs), quantifying all pairwise similarities of all patterns. These RSMs can then be compared to other RSMs based on other data.

In this study, we used RSA for two purposes. First, across participants we directly compared the time courses of MEG RSMs evoked by the *same* exemplar with MEG RSMs evoked by *different* exemplars. This allows an estimate of the generalizability of representations across exemplars and thus the extent to which a representation reflects high-level versus low-level properties, assuming that a generalized representation indicates a more high-level, conceptual representation. Second, we used RSA to study the relationship between evoked MEG activity patterns and computational, semantic, and behavioral models. In particular, we wanted to identify time periods at which the MEG responses reflected predominantly behavioral judgments, which we take as an index of high-level conceptual processing. To do this, we quantified the unique and shared variance of each model RSM with RSMs based on MEG activity patterns.

#### Construction of MEG similarity matrices

MEG RSMs were constructed as follows. For each time point, we averaged the preprocessed MEG data for all 36 trials of each object concept, yielding 84 object concept MEG patterns. Then we computed the similarity between all pairs of those 84 patterns across *P* principal components using a Spearman correlation, yielding an 84 × 84 MEG RSM for each time point. We then analyzed these RSMs further for the two purposes described above.

#### Comparison of low-level image similarity between image sets

To quantify the low-level similarity between image sets directly, we computed the pixelwise similarity across both image sets, concatenating the three color channels of each images to a vector and calculating the Spearman correlation between image vectors. This resulted in an 84 × 84 matrix, with the diagonal corresponding to the similarity within each object concept across image sets (e.g. “baby” in Image Set 1 with “baby” in Image Set 2) and the off-diagonals corresponding to the similarity across object concepts across object concepts (e.g. “baby” in Image Set 1 with “woman” in Image Set 2). In addition, as a computational model of low-level object processing we computed the GIST features (Oliva & Torralba, 2001) for each object image using default model parameters. We then calculated the Spearman correlation between those feature vectors in the same manner as described for pixelwise similarities.

#### Generalization of MEG similarity patterns across exemplars

To determine time periods that generalize between representations of object exemplars, we compared the time courses of similarity of RSMs *within* each image set to the similarity *between* image sets (see e.g. Guggenmos et al., 2018, for a similar methodological approach). To this end, we split data between the groups for Image Set 1 and Image Set 2 and conducted within- and between-group split-half correlation analyses with the RSMs for each participant. We chose a repeated subsampling procedure within group to allow us to use the same analysis within and between groups. The following analyses are described for one RSM at one time point, but were repeated for all time points.

Within each group of participants (*n* = 16), we randomly assigned participants’ RSMs to one of two arbitrary subsets of 8 participants and averaged participants’ RSMs within subsets. Next, we calculated the Spearman rank correlation coefficient between the lower triangular part of each 84 × 84 matrix, separately for every time point. We repeated this split-half analysis 1000 times with novel assignments of participants and averaged across repetitions, yielding a time course of within-exemplar correlation. The same procedure was completed for the between-group split-half analysis, but here the two subsets were each drawn from eight randomly selected participants in each group, yielding a time-course of between-exemplar correlations.

To assess statistical significance, we conducted a randomization test. We repeated the analysis above 1000 times (i.e. a total of 10^6^ split-half analyses, for both within-exemplar and between-exemplar comparisons). For each of those 1000 randomizations, we randomly permuted the rows and columns of the matrices in one of the subgroups before calculating Spearman’s *r*. *P*-values were determined as one minus the percentile of the original split-half analysis, and FDR-corrected to *p* < 0.05.

#### Representational similarity matrices for computational models and behavior

To identify and characterize the temporal evolution of the representational content of MEG responses in relation to behavior judgments, we chose multiple behavioral and computational models that we later compared to MEG data: a behavioral model based on the group mean behavioral similarity, a semantic model to capture similarity at the semantic level, and two layers of a deep neural network to capture different visual processing stages (low-to-mid level and high-level, respectively). The purpose of including those models was to identify the contribution of those processing stages to behavior, in order to gain a better understanding of the nature of the behavioral judgments. For a first comparison, we characterized the pairwise similarity of these models to assess their general similarity irrespective of MEG. We calculated Spearman’s *r* for each pair of models. Significance of correlations was tested using a randomization test: The rows and columns of one model RSM were randomly permuted before computing the Spearman’s *r* between with the other model RSM. This procedure was repeated 1,000 times to generate a null distribution of correlation coefficients, and results were deemed significant if they showed a higher correlation coefficient than the distribution cut-off determined by a level of *p* < 0.05.

##### Behavior

We generated an RSM for behavioral judgments by extracting the 84 × 84 distance matrices from each participant within a group and averaging them together. Next, we converted this distance matrix to an RSM by subtracting the distances from 1. Note that subsequent analyses only use the ranks of the entries of the distance matrices, which are simply inverted by this subtraction procedure. This step yielded two group-level behavior RSMs corresponding to Image Set 1 and Image Set 2.

##### Semantic model: Global Vectors for Word Representation (GloVe)

Global Vectors for Word Representations (GloVe) is an unsupervised algorithm that is trained on corpus word co-occurrence statistics to yield vector representations for words in the corpus, representing semantic relationships between words (Pennington et al., 2014). As a distributional measure of the semantic relatedness of words based on their shared linguistic contexts, GloVe is similar to other traditional co-occurrence models of word meaning but is particularly well-suited to the analysis here because of the high-dimensional similarity structure that shows semantic similarity between pairs of individual words, outperforming similar models in similarity tasks. As such, the structure of GloVe provides a fine-grained metric to evaluate how the representational space of MEG signals reflects semantic relationships as derived from shared lexical contexts. We chose 50-dimensional word vectors pre-trained on a 6-billion token Wikipedia database, extracted them for each object concept in the stimulus set and calculated Spearman’s *r* between each pair of vectors, generating an 84 × 84 RSM.

##### Visual model: Deep neural network VGG-F

We used the MatConvNet toolbox (Vedaldi & Lenc, 2015) to implement a pre-trained version of the Visual Geometry Group-Fast deep neural network (VGG-F DNN) (Chatfield et al., 2014) that was trained to perform the ImageNet ILSVRC 2012 object classification task. This network was chosen based on its high classification performance, ease of implementation, and suitability for our visual object concept stimuli. DNN representations for each image in both image sets were extracted from both convolutional layers (1-5) and fully-connected layers (6-8) of the network. We focused on representative examples of the convolutional and fully connected layers (3 and 7, respectively) to reflect low-to-midlevel vision and high-level vision, respectively. Within each layer, we calculated Spearman’s *r* between each of the object conditions that yielded an 84 × 84 RSM for both layers within each participant group. This yielded four distinct RSMs: DNN Layer 3 and Layer 7 for Image Set 1, and DNN Layer 3 and Layer 7 for Image Set 2.

#### Representational similarity analysis: Model comparisons to MEG

To directly compare each model to MEG activity patterns, we calculated Spearman’s *r* between the lower diagonals of the model variables and MEG RSMs at each time point within each group. These group-specific correlations were averaged together to yield a time course showing the level of correlation between the model and MEG responses. Upper and lower bounds for noise ceilings were determined within each of the two groups of participants according to Nili et al. (2014): The upper bound was estimated by calculating the correlation between each participant’s RSM and the mean group RSM *including* that participant, while the lower bound was estimated by calculating the correlation between each participant’s RSM and the mean group RSM *excluding* that participant. The upper and lower bounds from each group were averaged together to yield a mean noise ceiling across all participants. The statistical significance of this suite of representational similarity analyses was determined using randomization tests as described above, permuting the rows and columns of a given model RSM (behavior, GloVe, DNN Layer 3, DNN Layer 7) and for each randomization computing correlation time courses with the original MEG RSMs. Correlations were deemed significant if they exceeded a correlation cut-off determined by a level of *p* < 0.05 (FDR-corrected).

#### Establishing the unique and shared contributions of individual models

To determine the unique and shared variance between models and MEG signals, we conducted multiple linear regression analyses using the behavior RSM, DNN Layer 3 RSM, and DNN Layer 7 RSM as regressors to predict MEG RSMs from these variables (see See Groen et al., 2012; Lescroart et al., 2015; Greene et al., 2016, 2018; Hebart et al., 2018 for similar approaches). Given the complexities of describing the unique and shared variance partitions of more than three regressors, we decided to exclude the GloVe model, which showed the weakest correlation with MEG. Post-hoc analyses running different versions of the variance partitioning analysis (replacing the behavior model by GloVe, and separately replacing the DNN Layer 7 model by GloVe) demonstrated that the GloVe model overall explained less MEG variance than behavior (Supplemental Figure S4), while explaining very little unique variance (Supplemental Figure S5). By conducting a series of different multiple regressions with different combinations of model variables, this approach allows us to determine not only the unique MEG variance explained by each model RSM individually, but also the variance shared between any combination of model RSMs. Before conducting variance partitioning analyses, we averaged the group-specific RSMs of both image sets for behavior and DNN models, which yielded very similar results as compared to calculating them separately and averaging results afterwards. We extracted the lower diagonal from the mean MEG RSM at each time point as dependent variables, and assigned each of the models as independent variables. In sum, 7 regression analyses were performed at each time point that each included different combinations of models as regressors: 1) ‘full’ regression, including all three models (DNN Layer 3, DNN Layer 7, behavior), (2-4) ‘combined-predictor’ regression, including all pairwise combinations of two models (DNN Layer 3 and behavior, DNN Layer 7 and behavior, DNN Layer 3 and DNN Layer 7), and (5-7) ‘single-predictor regression’ including each model on its own. Subtracting the explained variance (*R^2^*) values of these different regression analyses yields portions of variance that are independently explained by each model, the variance that each model shares with the other two models, and the variance shared by all three.

For example, the unique variance explained by behavior (region c in the Venn diagram depicted in Figure 6a) is computed as the difference in *R^2^* between the full regression model (which includes all three regressors and therefore encompasses all regions described by the red, green and blue circle, i.e. a+b+c+ab+ac+bc+abc) and a regression model including only DNN Layer 3 and 7 (encompassing all regions described by the green and blue circle, i.e. a+b+ab+ac+bc+abc). Once the three regions of unique variance (a, b, and c) are obtained in this way, shared variances can be computed. For example, the variance shared by behavior and DNN Layer 7 (region bc) is computed by taking the *R^2^* resulting from including behavior and the third model (DNN Layer 3) (corresponding to all regions covered by the red and blue circle, i.e. a+ac+ab+abc+c+bc) and subtracting both the *R^2^* obtained when including DNN Layer 3 alone (blue circle, a+ab+ac+abc) as well as the unique variance explained by behavior (region c). Finally, the variance shared by all three models (region abc) is computed as the difference between the full regression *R^2^* and all the sum of all unique variances (a+b+c) and shared variances between all combinations of two models (ab+ac+bc). Statistical significance was determined using a randomization test as described above, randomizing columns and rows of model matrices 1000 times and repeating the original analysis. For a given iteration, the same randomization was used across all models to fulfill the assumptions of the randomization test. Significance cutoffs for *R^2^* were set to *p* < 0.05 (FDR-corrected).

## Results

Our aim in this study was to characterize the emergence of conceptual representations for visual objects. We applied multivariate decoding and representational similarity analysis to MEG data to examine (1) how object representations generalize across time and object exemplars, and (2) to elucidate the unique and shared contributions of behavioral judgments to measured MEG responses. The resulting temporal profiles inform us about stages of object processing from low-level visual to conceptual representations.

### Time-resolved representation of object identity

To characterize the time course by which neural signals in the human brain convey information about object identity, we used time-resolved multivariate decoding, conducting pairwise classification between MEG patterns in response to object stimuli (Figure 2a). Object identity information rose rapidly in response to stimulus presentation, with decoding accuracy peaking at 100 ms (mean accuracy: 91.1 %), followed by a slow decay of information that remained significantly above chance after stimulus offset and for the duration of the trial time window (1000 ms post stimulus onset). These results indicate that we were able to detect the temporal unfolding of object-identity information encoded in MEG signals with high accuracy, establishing a correspondence to previous research demonstrating that discriminable object representations emerge well within 100 ms of visual recognition (Carlson et al., 2013; Cichy et al., 2014). Further, these results lay an important foundation for the following analyses in which we delineate what information specifically contributes to these discriminable object representations.

**Figure 2.**
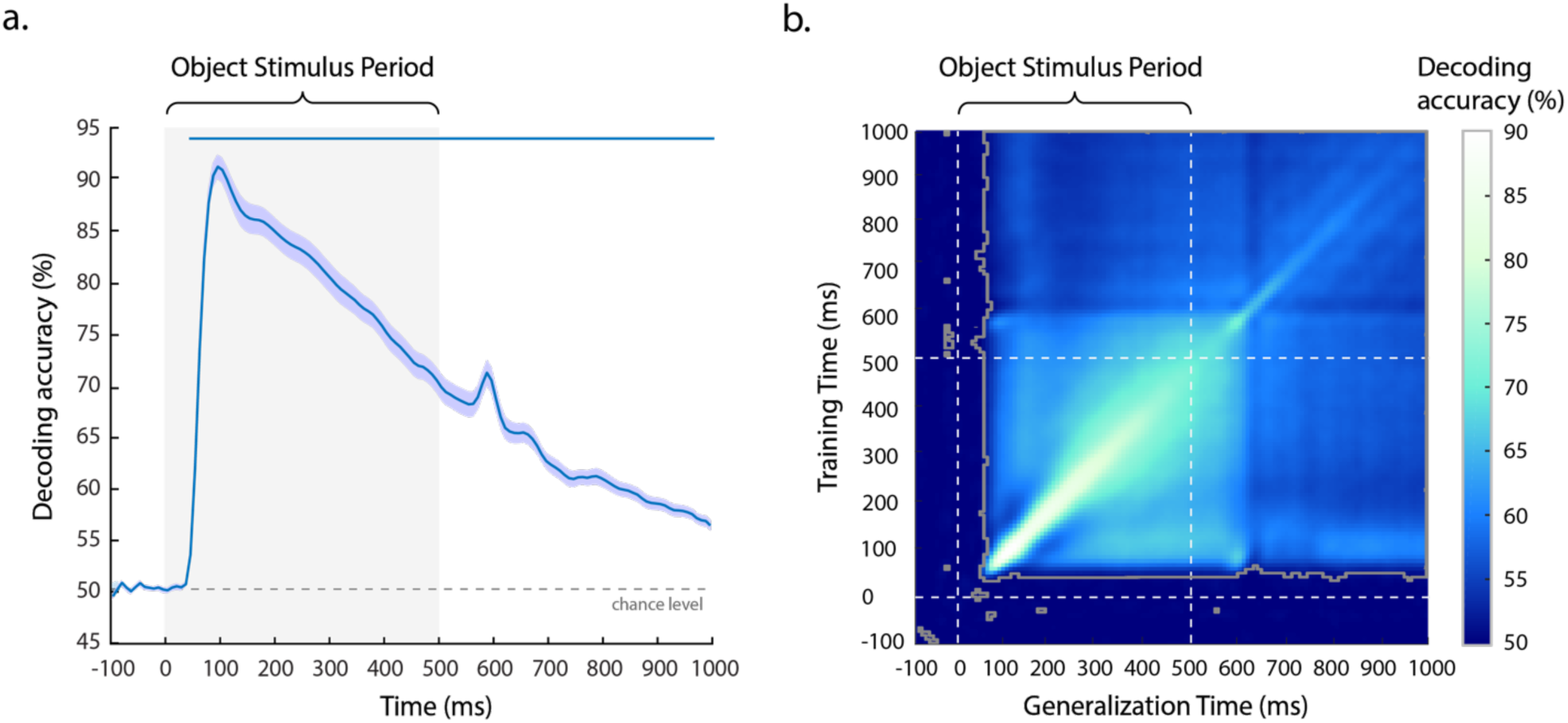
**a.** Time-resolved multivariate decoding of object identity across the trial. After onset of the object stimulus (Object Stimulus Period), pairwise object decoding accuracy increased rapidly, followed by a slow decay towards chance over the duration of the trial. Error bars reflect SEM across participants for each time point separately. Significance is indicated by colored lines above the plot (non-parametric cluster-correction at *p* < 0.05). **b.** Temporal generalization matrix for object identity. The y-axis depicts the classifier training time relative to stimulus onset, and the x-axis classifier generalization time relative to stimulus onset. Dotted lines indicate stimulus onset and offset. Areas bounded by a grey line contain significant temporal cross-decoding accuracy values (*p* < 0.05, FDR corrected). See Supplemental Figure S2, for horizontal cross-sections across the temporal generalization plots.

### Temporal generalization of object information

While time-resolved multivariate decoding reveals the temporal evolution of discriminable object representations, it does not inform about the dynamics and stability of those representations across time. To identify the degree to which object representations generalize across time, we ran a temporal generalization analysis by training a classifier on data at every time point and testing it at all other time points. This yielded a temporal generalization matrix (Figure 2b), with the diagonal representing training and testing at the same time points, mirroring the results presented in Figure 2a. In a temporal generalization matrix, a dynamic representation would be characterized by high accuracies around the diagonal and low accuracies everywhere else, indicating little generalization across time. In contrast, a stable neural representation would exhibit high decoding around the diagonal but also in the off-diagonal time points, demonstrating a similar representation across time.

Our results exhibited significant generalization from ~70 ms onward, demonstrating a shared representational format across the entire trial. While this result reveals a persistent representation across time, the strength of generalization varies. Focusing on the first half of the stimulus presentation period, the results revealed a period of increased temporal dynamics between ~70-250 ms, indicated by the high decoding accuracy on the diagonal and lower decoding accuracies away from the diagonal. This result suggests a relatively dynamic representational format in this phase of visual processing. After ~250 ms, there was increased generalization away from the diagonal, indicating a more persistent, shared representational format during this later phase of visual processing. Interestingly, there was a generalization period between time windows of ~70-100 ms and ~250-550 ms, suggesting an overlap of representations between early visual and later conceptual processing. The markedly lower information generalization between 150-250 ms and all other time points suggests the information dynamics at these points are computationally dissimilar from other stages of processing.

Taken together, these results reveal relatively weak but significant persistence of stable object information throughout the entire trial. On top of this, the results reveal a general broadening of information generalization after an early phase of visual processing. While the results of this temporal generalization analysis do not reveal multiple distinct stages of processing, this broadening suggests early dynamic neural activity followed by the emergence of more stable object representations around 250 ms.

### Criterion I for conceptual object representation: Generalization between object exemplars

Having established the time course of object identity-specific information, we investigated when those brain responses reflect conceptual object representations. One prerequisite of a conceptual object representation is a similar representational format between multiple exemplars of the same object, since a conceptual representation is expected to generalize beyond each individual exemplar. The data collected from Image Set 1 and 2 allow direct comparison of representational similarity across exemplars for the same visual object concept (Figure 3). We expected some low-level features to be shared across object exemplars, but that this tendency would be reduced as compared to the same exemplar. Indeed, when comparing the low-level similarity between exemplars of image sets, the similarity was higher within object concept than between object concepts (mean pixelwise similarity within: *r* = 0.15, between: *r* = 0.04; mean GIST similarity within: *r* = 0.58, between: *r* = 0.41), demonstrating some preserved similarity between object exemplars. However, the overall similarity was strongly reduced, and the maximal similarity across image sets was for the same object concept in only 19% of the cases (based on the GIST similarity), demonstrating a strongly reduced low-level similarity between image sets.

**Figure 3.**
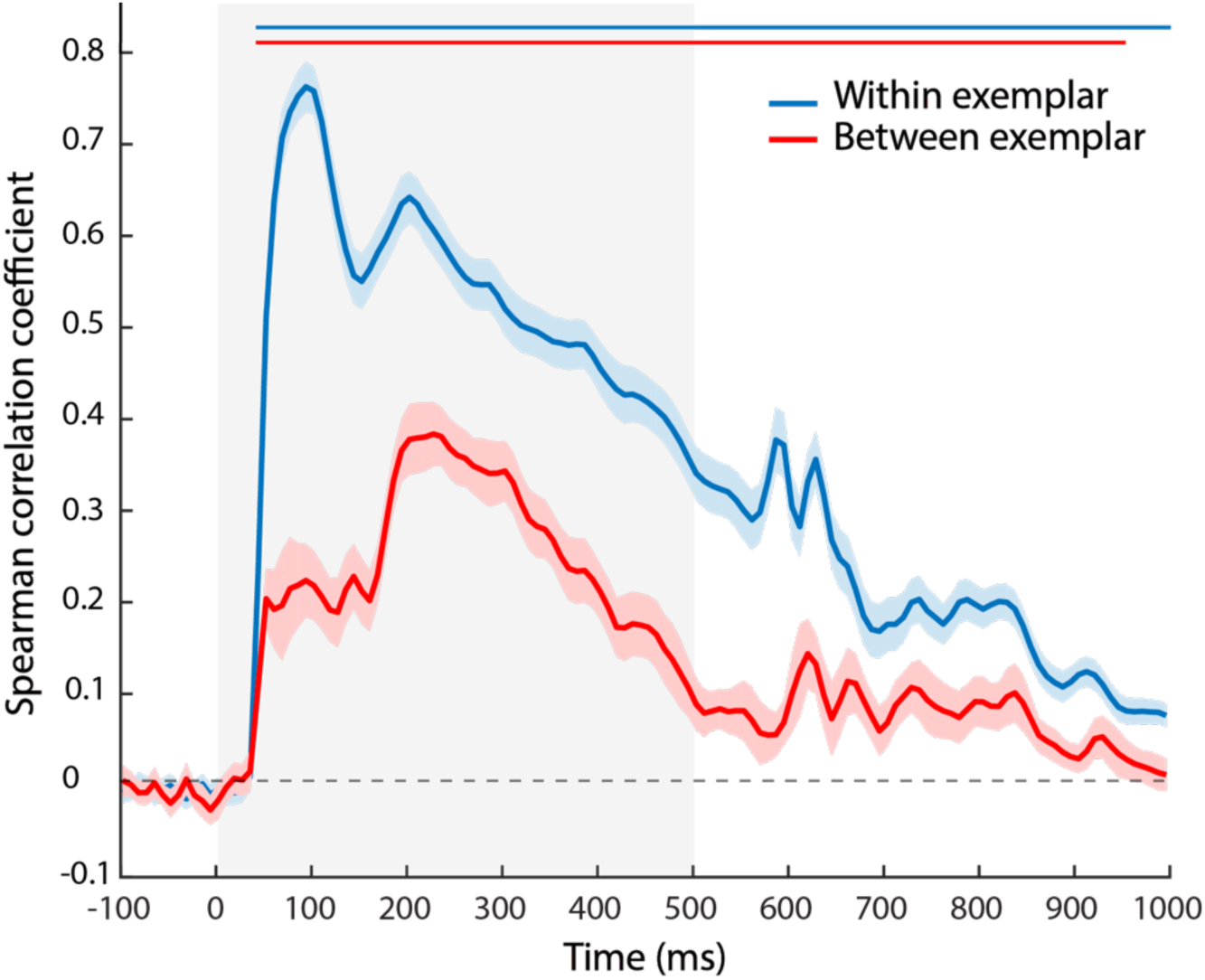
Within and between exemplar correlation of MEG RSMs. Within-exemplar correlation was generally higher than between-exemplar correlation. Both within and between-exemplar correlations revealed an early peak (93 ms) and a late peak (202 and 227 ms, respectively), with the early peak being higher than the late peak for within-exemplar correlations, and the late peak being higher than the early peak for between-exemplar correlations. Error bars reflect SEM. Significance is indicated by colored lines above the accuracy plot (non-parametric cluster-correction at *p* < 0.05).

Having demonstrated the reduction in low-level similarity between image sets, we measured this generalization of object concept-specific information by (1) calculating the correlation of within-exemplar MEG RSMs for participants who were shown the same object exemplar and (2) calculating the generalization of between-exemplar MEG RSMs for participants who were shown different object exemplars. Then we compared the shape of these MEG correlation time courses.

A comparison of within-exemplar and between-exemplar MEG RSM correlations revealed a generally higher correlation within-exemplar than between-exemplar (mean difference across time: Spearman’s *r*: 0.18, *p* < 0.001, randomization test), indicating that differences between exemplars persisted throughout most of the trial. Reliable structure for within-exemplar MEG RSMs emerged rapidly, peaking at 93 ms (mean Spearman’s *r*: 0.77). This was followed by a fast drop in correlation, and then another rise beginning around 160 ms and peaking at 202 ms (mean Spearman’s *r*: 0.65), after which within-exemplar correlations decreased steadily for the duration of the trial while remaining significantly above chance. The correlation of between-exemplar MEG RSMs also initially increased rapidly, but then reached a plateau at a comparably low level of correlation between ~70 and ~160 ms (mean Spearman’s *r*: 0.21). Importantly, between-exemplar reliability then increased again after ~160 ms, peaking at 227 ms (mean Spearman’s *r:* 0.39). Between-exemplar correlation then slowly decayed back to 0, but remained significantly above chance until 960 ms after stimulus presentation.

These results reveal an important dissociation: While within-exemplar correlations reached their maximum around 100 ms, between-exemplar generalization was maximal around 200 ms. Thus, this analysis reveals an early processing stage during which generalization is limited by the variable visual features of each individual exemplar, and a later processing stage where the increased generalization likely reflects the development of a more conceptual object representation that is consistent across exemplars.

### Comparison of behavior and computational models of low-level and high-level processing

To quantify how the RSMs derived from behavior (perceptual judgments, visualized in Figure 4b), GloVe (lexical semantics), DNN Layer 3 (low/mid-level visual information), and DNN Layer 7 (high-level visual information) relate to one another, we computed the correlation between each pair of model RSMs (Figure 4a). For visualization purposes, we applied hierarchical clustering to independent pilot data of the behavioral task to sort objects depicted in the model RSMs (Figure 4a). All model correlations were significant at a level of *p* < 0.001 (randomization test). An estimate of the upper noise ceiling for possible model correlation values was calculated by the correlation between behavior RSMs for the two groups of participants (Spearman’s *r* = 0.64). The greatest similarity to behavior was shown by the GloVe model. There was low similarity of convolutional DNN Layer 3 with behavior and GloVe, but much greater similarity for fully-connected DNN Layer 7. These results suggest an increase of semantic, behaviorally-related information contained in the representational structure of the DNN Layer 7 as compared to Layer 3. Note that the lowest correlation observed was between DNN Layer 3 and behavior RSMs, indicating that behavior was not strongly driven by low-to mid-level responses. As a post-hoc analysis, we added the comparison of behavior to the GIST RSM, which was even lower (Spearman’s *r* = 0.02), highlighting the low explanatory power of low-level features in behavioral judgments in the present study.

**Figure 4:**
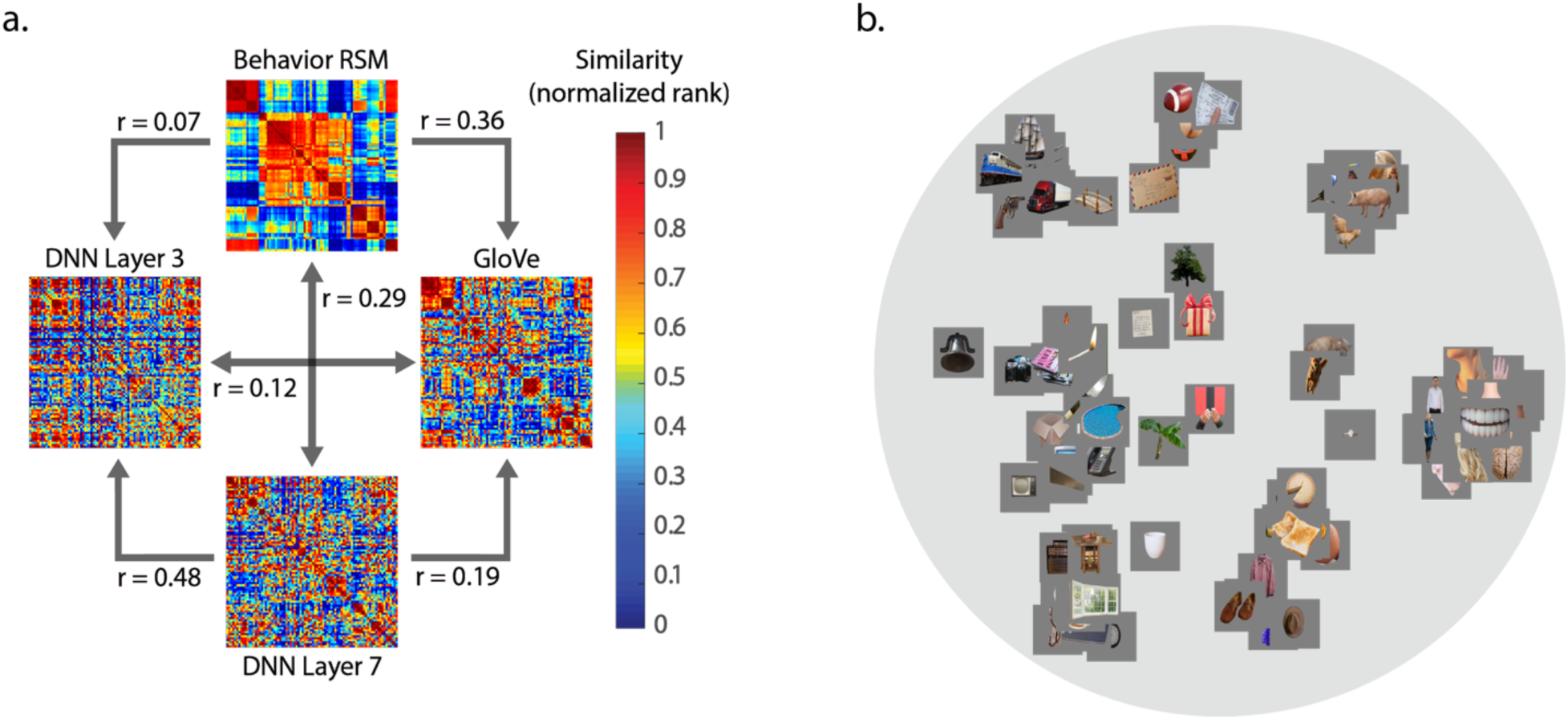
**a.** Explicit comparison of computational models and behavior using RSA. Models compared are group average behavior, GloVe, DNN Layer 3 and DNN Layer 7. RSAs are plotted as ranks for higher visual contrast. Objects are sorted based on clustering generated from independent pilot data. **b.** Group average inverse MDS plot generated from behavioral arrangement task.

### Criterion II for conceptual object representation: Behavioral and computational modeling of MEG data

To determine when there is a relationship between the MEG signal and high-level behavioral judgments, satisfying Criterion II, we first evaluated the time course of similarity between behavioral judgments and the MEG activity patterns (Figure 5). Further, to establish whether this relationship was uniquely explained by behavior, we additionally compared MEG to the computational models described above. Every model tested exhibited significant correlations with MEG activity patterns within the first 200 ms of visual processing. DNN Layer 3 showed peak correlation with MEG at 118 ms after stimulus onset (Spearman’s *r* = 0.33), while DNN Layer 7 showed peak correlation with MEG at 151 ms (Spearman’s *r* = 0.23). Further, the GloVe model was most strongly correlated with MEG at 151 ms (Spearman’s *r* = 0.13), and behavior at 160 ms (Spearman’s *r* = 0.16). Additional within-subject analyses, i.e. comparing each individual’s behavioral RSM to their MEG RSM, revealed a very similar pattern of results but lower overall correlations (peak Spearman’s *r* = 0.06) and no significant benefit of within-subject over between-subject analyses matched in size (all *p* > 0.12).

**Figure 5.**
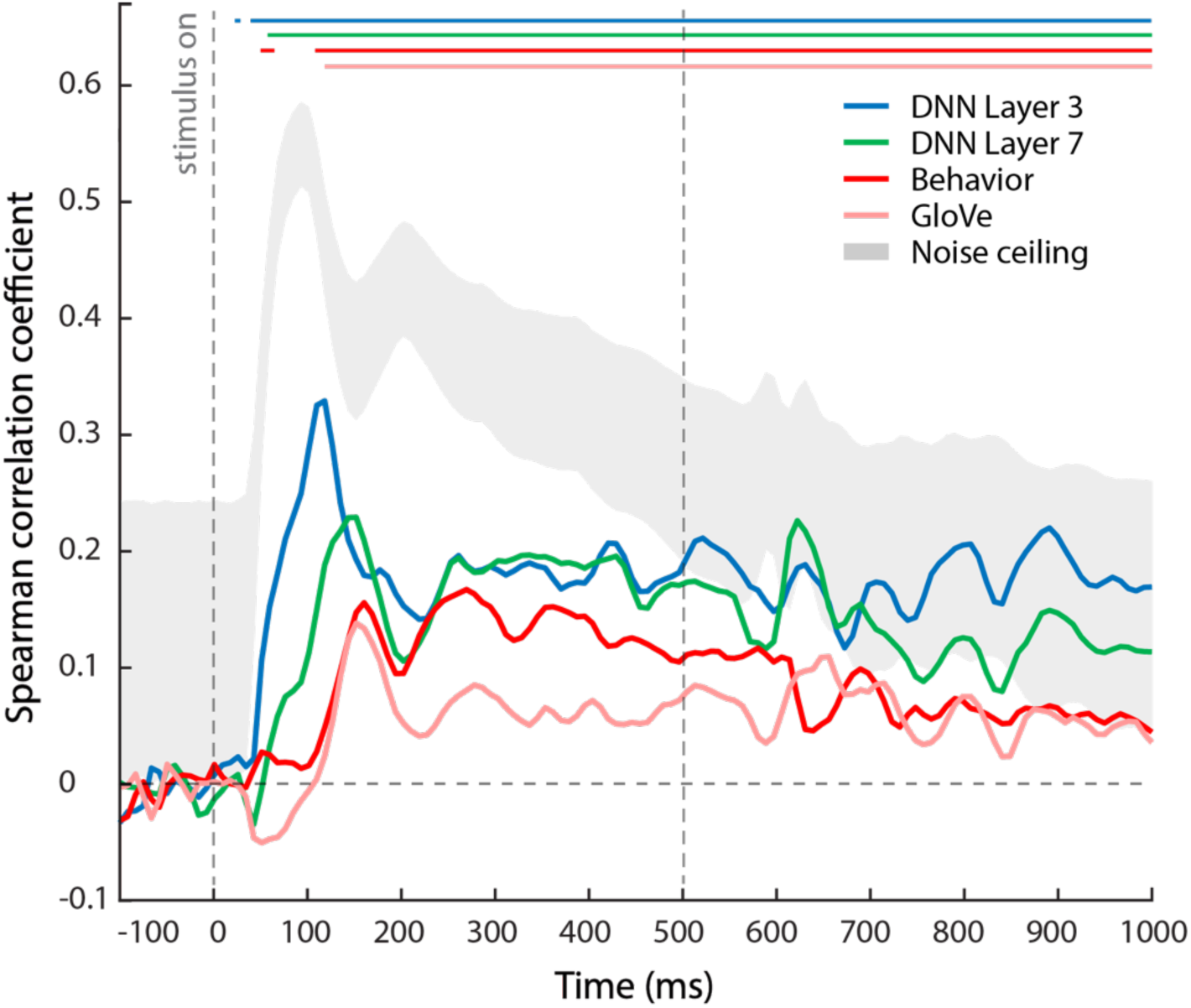
Results of model-based representational similarity analysis with MEG data. Comparison includes models based on DNN Layer 3, DNN Layer 7, GloVe and behavior. The results exhibit a progression of peaks from DNN Layer 3 to behavior, suggesting a temporal evolution of the underlying representation from more low-level to higher-level/conceptual. Grey shaded area depicts the noise ceiling. Significant time points are indicated by a colored line above the plots (*p* < 0.05, FDR-corrected permutation test).

This sequence of peaks suggests an evolution from low-level visual to high-level conceptual representations, with the relationship to behavior peaking latest in time. However, given the significant correlations of all models with MEG throughout most of the trial and the presence of significant correlation between the models themselves (Figure 4a), it is unclear to what extent a given correlation was uniquely explained by one model, or whether this correlation could equally well be explained by other models. For example, the correlation of both DNN Layer 7 and behavior with MEG signals after 150 ms raises the question whether the behavioral correlations can be fully explained by the features represented in the DNN models.

### Variance Partitioning: Shared and unique model contributions

To provide a deeper understanding of the unique contributions of different models to MEG variance and how much explanatory power they share with behavioral judgments in explaining MEG variance, we conducted a variance partitioning analysis in which we compared the results of different multiple regression analyses applied to MEG RSMs (see Methods; Figure 6a). We first considered the total percent of variance in the MEG RSMs explained when all three predictors are combined in a single regression model (‘full model’) in comparison to the percent variance explained by each model separately (Figure 6b). Since variance explained by each model separately is identical to the square of the model correlation, the results of this analysis are very similar to those of the previous section presented in Figure 5, with the only difference that these results were collapsed across groups before conducting variance partitioning. Explained variance of DNN Layer 3 peaked at 118 ms (*R*^2^: 11.0 %), DNN Layer 7 at 151 ms (*R*^2^: 7.0 %), and behavior at 160 ms (*R*^2^: 4.8 %). Importantly, however, the dashed line indicates how these contributions relate to the total variance accounted for by all three models combined. At its peak at 118 ms, the full model explains 11.6 % of the variance, which is similar to the amount of variance explained by DNN Layer 3 alone, suggesting that all variance captured at this time-point can be attributed uniquely to DNN Layer 3, with limited additional contribution of DNN Layer 7 or behavior. At later time points, however, the full model always substantially explains more variance than the individual predictors, providing a first clue that some or all of these predictors contribute unique (i.e., additive) variance.

**Figure 6.**
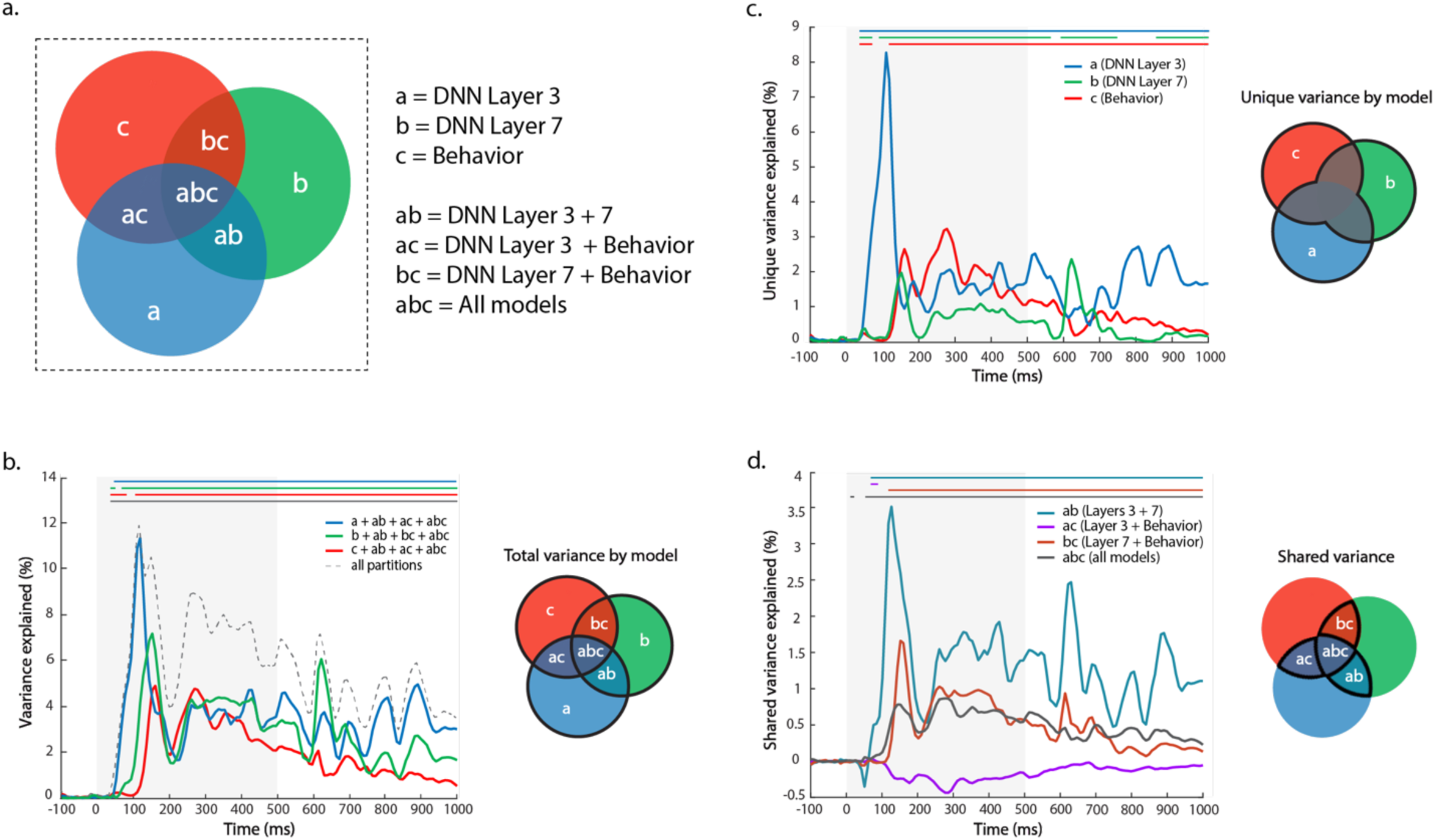
Time-resolved variance partitioning: Total, shared, and unique MEG variance explained by models: DNN Layer 3, DNN Layer 7, and behavior. **a.** Schematic of unique and shared variance components using a Venn diagram. **b.** Percent MEG variance explained by each model independently (colored lines), and total MEG variance explained at all time points (dotted line). **c.** Unique variance explained by each model. **d.** Shared variance between different model combinations. Significant time points are indicated by a colored line above the plots (*p* < 0.05, FDR-corrected permutation test).

To directly quantify the unique and shared variance of each model, we compared the regression outcomes with different model variables included (Figures 6c, 6d). The unique MEG variance explained by DNN Layer 3 peaked very early in time, at 109 ms (*R*^2^: 8.3%). DNN Layer 7 peaked next at 151 ms (*R*^2^: 2.0 %), followed closely by behavior at 160 ms (*R*^2^: 2.6 %), with a second peak at 277 ms (*R*^2^: 3.2 %). Importantly, DNN Layer 3 explained the most unique variance until 143 ms, after which behavior predicted the most unique variance until ~400 ms. Thus, while all three models (DNN Layer 3, DNN Layer 7 and behavior) captured some unique variance in MEG activity throughout the trial, behavior dominated after around 150 ms.

Finally, to complete the picture, we partitioned the variance into shared contributions from combinations of the different models. Both DNN Layers contributed the most shared variance across all time points after stimulus onset, which is perhaps not surprising considering that both layers are derived from the same computational model. This shared variance between DNN Layer 3 and Layer 7 peaked at 126 ms (*R*^2^: 3.5 %). Interestingly, the shared variance between behavior and DNN Layer 7 demonstrated a clear peak at 151 ms (*R*^2^: 1.7 %), suggesting that it is around this time-point that DNN Layer 7 best captures neural information that is also reflected in behavior. The shared variance between DNN Layer 3 and behavior was slightly negative, a result that is not untypical for variance partitioning, indicative of small suppression effects (Pedhazur, 1997) and suggesting that DNN Layer 3 does not capture information that is relevant for behavioral judgments.

It is possible that DNN Layer 3 did not accurately capture the low-level responses. For this reason, we ran additional variance partitioning analyses, replacing DNN Layer 3 with the GIST model. The GIST RSM exhibited a strong correlation with the DNN Layer 3 RSM (Spearman’s *r:* 0.65). As expected from this correlation, the variance partitioning results were qualitatively very similar (Supplemental Figure S3), demonstrating that DNN Layer 3 likely captured relevant low-level responses.

Collectively, the variance partitioning results indicate that behavioral judgments are reflected in the MEG response above and beyond what is captured by the DNN, with behavioral judgments explaining the most unique variance between 200 and 400 ms after stimulus onset. Further, before 150 ms, DNN Layer 3 explains the most variance, suggesting that representations prior to this point are unlikely to be conceptual in nature.

## Discussion

In this study, we investigated the temporal evolution of visual object representations. In particular we focused on determining a lower bound for the emergence of conceptual representations of objects. We proposed two criteria that would reflect conceptual representations: 1) generalization of representations between different exemplars of the same object, and 2) relationship to high-level behavioral judgments. We find qualitatively different processing of objects over time: Early responses (< 150 ms) were characterized by exemplar-level representations and similarity with computational visual models, whereas later responses (> 150 ms) showed increasing generalization across exemplars and similarity with behavioral judgments, with greater stability of representations over time.

To evaluate generalization of representations reflecting conceptual processing, we compared the representational structure of MEG responses, both within exemplar and between sets of exemplars. This analysis revealed two interesting features. First, between-exemplar generalization was found to be consistently lower than within-exemplar generalization, demonstrating the persistence of exemplar-specific responses. This reduced between-exemplar generalization likely reflects the impact of low-level features varying between different exemplars. The fact that this advantage is maintained throughout the trial, suggests some persistence of low-level feature representation. This interpretation is supported by the temporal generalization even for very early time points and the variance explained by DNN Layer 3 (which likely corresponds to early to mid-level visual processing, Cichy et al., 2016a; Güçlü & van Gerven, 2015; Wen et al., 2017), throughout the trial. Second, both within and between-exemplar generalization showed two distinct peaks, one early around 100 ms, and another late around 200 ms. However, their relative amplitude was reversed: While the early peak was stronger than the second within-exemplar, this pattern was reversed between-exemplar. This striking increase in generalization between exemplars that occurs for the later peak suggests the emergence of a common representation across exemplars, a key marker for conceptual representations. Together these results suggest that the earliest time point for the emergence of conceptual representations is around 150 ms, but also suggest a prolonged representation of low-level visual features.

To evaluate the relationship to high-level behavioral judgments, we compared models derived from behavior, semantics (GloVe), and computational vision (DNN) with the MEG response to objects. We found that all models show significant correlation with the MEG response throughout most of the trial. The early DNN layer showed the strongest and earliest correlation, while the GloVe model showed the weakest correlation. This result highlights the importance of testing multiple models rather than relying on a significant effect for a single model. Since the models themselves are correlated (Figure 4), this demonstrates that testing multiple models is *also* not sufficient; it is important to determine the unique and shared variance explained by the different models (Lescroart et al., 2015; Groen et al., 2012; Greene et al., 2016; Hebart et al., 2018; Groen et al., 2018), motivating our variance partitioning analysis. Given the complexities of describing the unique and shared variance partitions of more than three model variables, we decided to exclude one of the four. Since the GloVe model showed the weakest correlation with MEG and was mostly subsumed by the behavioral model, we focused on the DNN and behavioral model variables.

The variance partitioning revealed several important features. Focusing on the unique contribution of each model variable, it becomes clear that DNN Layer 3 dominates early MEG responses peaking at 100 ms, whereas behavior explains the most variance after 150 ms, peaking at 270 ms. This result fulfills our second criterion – relationship with high-level behavioral judgments – converging with the results of both the temporal generalization analysis and the representational generalization across exemplars in identifying the time period after around 150 ms as reflecting a lower bound for the emergence of conceptual representations. Focusing on the shared contribution of model variables, the results largely reflect the correlations between model variables (Figure 4), e.g. no shared variance between DNN Layer 3 and behavior, high shared variance between Layers 3 and 7 of the DNN model. However, they provide important information about the timing of the shared variances. In particular, the shared variance between DNN Layers 3 and 7 persisted even late in time, again suggesting a sustained representation of low-level visual information.

Our results are generally consistent with prior work investigating how visual processing of objects evolves over time, showing the gradual emergence of high-level representations (Contini et al., 2017). While early signals reflect low-level visual features (e.g. Groen et al., 2013; Cichy et al., 2014; Coggan et al., 2016), later signals reflect perceptual similarity (Wardle et al., 2016), some tolerance for changes in size and position (Isik et al., 2014), categorical processing (Carlson et al., 2013; Cichy et al., 2014), and correlate with task performance and reaction times (Van Rullen and Thorpe, 2001; Philiastides and Sajda, 2006; Martinovic et al., 2008; Ritchie et al., 2015). Further, comparisons of deep neural networks with MEG have revealed a correspondence of early layers with earlier MEG responses, likely reflecting initial stages of processing in early visual cortex, while higher layers reflect later stages of processing in occipitotemporal cortex (Cichy et al., 2016b; Seeliger et al., 2017). Our results significantly extend these results by establishing a lower bound for the development of conceptual representations.

Other studies have also investigated high-level conceptual processing over time using explicit semantic feature models (Clarke & Tyler, 2015) or behavioral judgments (Cichy et al., 2017). For example, Clarke and colleagues showed semantic feature effects before 120 ms, although including basic visual features based on the HMAX model revealed unique semantic contributions to MEG signals only after ~200 ms (Clarke et al., 2013; Clarke et al., 2014). In contrast to these studies, we used more recent deep convolutional neural networks which have been shown to be more closely tied to neural and behavioral data (Khaligh-Razavi et al., 2016; Jozwik et al., 2017; Cichy et al., 2016a). Further, we operationalized high-level conceptual processing, using both a computational semantic model based on semantic co-occurrence statistics (GloVe model), as well as behavioral judgments of object similarity that we take to more broadly reflect conceptual processing. Indeed, our results suggest that MEG variance explained by the GloVe model was comparably low and mostly covaried with behavioral judgments, suggesting that conceptual representations extend beyond those relationships captured by the GloVe model. Despite these differences, our results are generally consistent with the results of Clarke and colleagues, but suggest a lower bound for conceptual processing around ~150 ms (see also Cichy et al., 2017). Further, we show that the computational visual model and behavioral judgments explain shared variance even prior to 150 ms. This shared variance indicates that the neural activity captured by compational models is behaviorally relevant and argues against a strong distinction between (low-level) visual features on the one hand, and high-level conceptual processing on the other. At the same time, the presence of significant unique variance explained by behavior after 150 ms suggests that not *all* aspects of conceptual object representations reflected in MEG activity are explained by current generations of computational visual models.

While our study provides insight into the development of conceptual representations, there are some important considerations. First, we used behavioral similarity judgments using the multi-arrangement task (Kriegeskorte & Mur, 2012) to index conceptual processing. However, this choice of method might constrain the ability to capture conceptual representations. While the behavioral judgments explain more variance in the MEG signal than the semantic GloVe model we tested, we do not know what aspects of conceptual processing are reflected in those judgments. Further, it is unclear how sensitive those behavioral judgments are to the context imposed by the stimuli and instructions. Second, we only employed two exemplars per object concept to test generalization of representations, and this may not have contained sufficient variability to fully disentangle low-level and high-level processing. In future work, it would be useful to carry out similar analyses while presenting multiple image sets to each participant, which would allow within-subject exploration of differences in temporal generalization across exemplars. Finally, in the future alternative similarity metrics for MEG-based RSA, such as the cross-validated Mahalanobis distance could be applied that may increase the robustness over the current approach (Guggenmos et al., 2018). Future studies should consider broader sets of stimuli, different behavioral tasks, and alternative computational models that may better match the MEG signal.

In conclusion, by focusing on two criteria for conceptual object representations we provide an estimate for a lower bound for the emergence of conceptual object representations of around 150 ms. Prior to this time, our results demonstrate limited generalization across object exemplars and time, and importantly little unique contributions of behavioral judgments to the MEG response. The multifaceted nature of our findings here show that the combination of neural data, behavior, and models are a viable method to probe the temporal dynamics of object recognition and allow us to establish a novel profile of emergent conceptual representations in time.

## Acknowledgements

This work was supported by the Intramural Research Program of the National Institute of Mental Health (ZIA-MH-002909) - National Institute of Mental Health Clinical Study Protocol 93-M-0170, NCT00001360, a Feodor-Lynen fellowship of the Humboldt Foundation to M.N.H., and a Rubicon Fellowship from the Netherlands Organization for Scientific Research to I.I.A.G.

